# Lineage-based scaling of germline intercellular bridges during oogenesis

**DOI:** 10.1101/2023.08.18.553876

**Authors:** Umayr Shaikh, Kathleen Sherlock, Julia Wilson, William Gilliland, Lindsay Lewellyn

## Abstract

The size of subcellular structures must be tightly controlled to maintain normal cell function. Despite its importance, few studies have determined how the size of organelles or other structures is maintained during development, when cells are growing, dividing, and rearranging. The developing egg chamber is a powerful model in which to study the relative growth rates of subcellular structures. The egg chamber contains a cluster of sixteen germline cells, which are connected through intercellular bridges called ring canals. As the egg chamber grows, the germline cells and the ring canals that connect them increase in size. Here, we demonstrate that ring canal size scaling is related to lineage; the largest, “first born” ring canals increase in size at a relatively slower rate than ring canals derived from subsequent mitotic divisions. This lineage-based scaling relationship is maintained even if directed transport is reduced, ring canal size is altered, or in egg chambers with twice as many germline cells. Analysis of lines that produce larger or smaller mature eggs reveals different strategies could be used to alter final egg size.

**Summary Statement:** Using the fruit fly egg chamber as a model, this study demonstrates that the size and scaling of germline intercellular bridges vary based on lineage.

## Introduction

A fundamental question in biology is how size is regulated. There is an extensive literature describing the relationship between body size and the size of organs or appendages (Gayon, 2000; Huxley and Teissier, 1936), but we are just beginning to understand how size is regulated at the cellular level. Cells vary in size from one organism to another and from one cell type to another, but an essential feature of all cells is that they must regulate the size and number of their subcellular structures. Many studies have described the scaling of various organelles with cell size, including the nucleus, the mitotic spindle, mitochondria, the ER, and others (Marshall, 2020), but very little is known about how subcellular scaling is established and maintained within developing tissues, where cells are growing, dividing, and rearranging.

The developing fruit fly egg chamber is a powerful model system to study scaling of subcellular structures. The egg chamber is the functional unit of oogenesis which will ultimately produce the mature egg; within the egg chamber, the developing oocyte and its supporting nurse cells are surrounded by a layer of epithelial cells. The oocyte is connected to the nurse cells through intercellular bridges, called ring canals, which provide a channel for the delivery of essential materials from the nurse cells to the oocyte. From stages 2-14, the egg chamber increases in volume by approximately 1,000-fold (Fig. 1A; Ali-Murthy et al., 2021); therefore, the egg chamber provides us with the opportunity to learn how the size of subcellular structures is coordinated during tissue growth.

**Figure 1:**
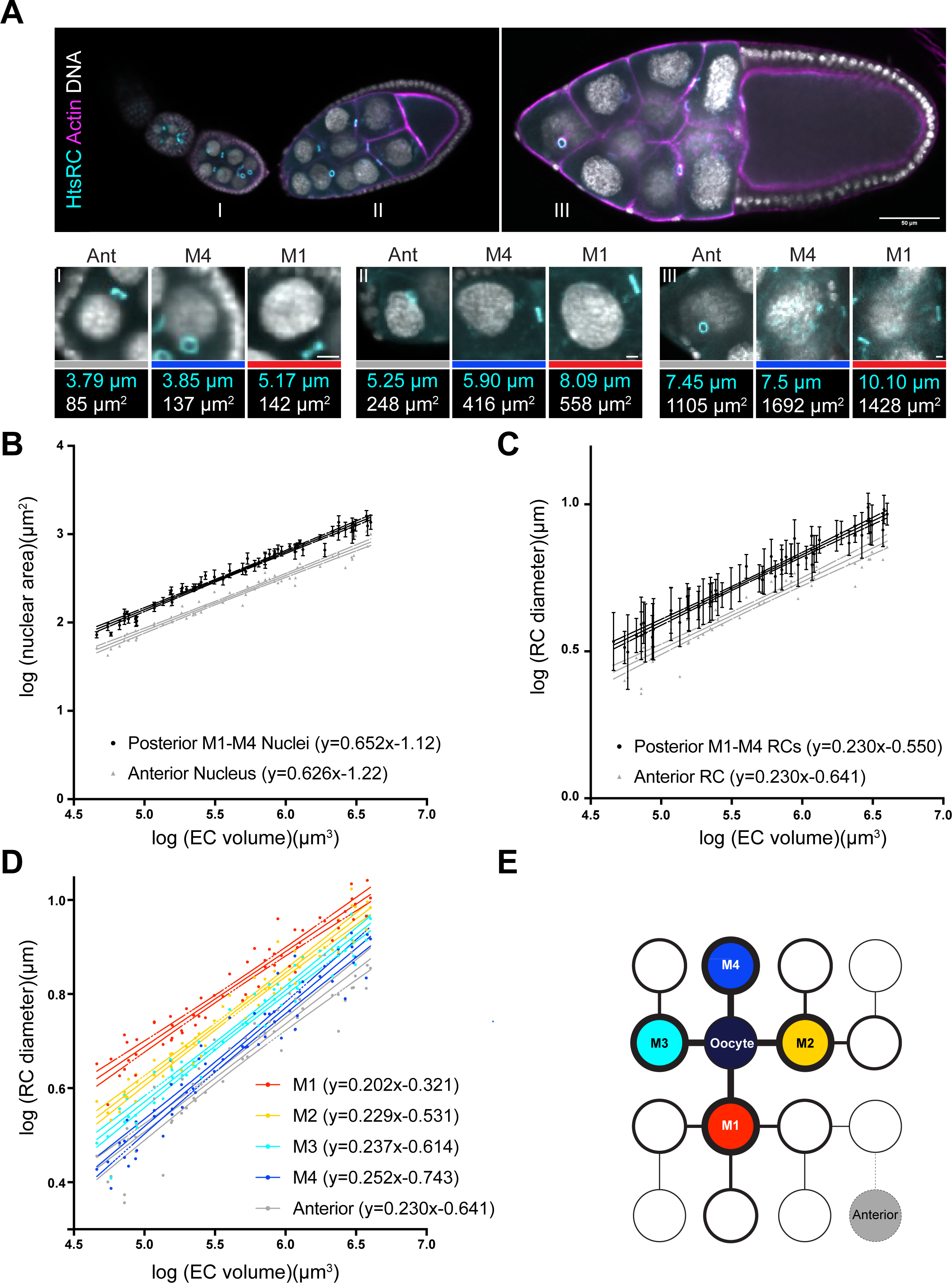
Ring canal size and nuclear size vary within the egg chamber. (A) Single plane confocal images of stage 5 (I), early stage 9 (II), and stage 10 (III) control egg chambers (*w^1118^/+;;matαTub-GAL4*) stained with an HtsRC antibody, phalloidin, and DAPI. Panels below show anterior, M4, and M1 ring canals from each of the indicated egg chambers (which were located in different z-planes than the one shown in the larger panels); ring canal identity is indicated by the colored bars, which match those used in (E). Measurements of these ring canals and associated nuclei are included below; measurements were made in the z-plane where each structure was in focus. Scale bar in larger panel is 50 µm; scalebars in smaller panels are 5 µm. (B-D) log-log plots of egg chamber (EC) volume, nuclear area, or ring canal (RC) diameter from stage 5-10b. All four posterior nuclei and ring canals (M1, M2, M3, and M4) were measured. Error bars in B,C are s.d. n=61 egg chambers. 95% confidence intervals are shown. (E) Cartoon showing the stereotypical connections that result from the four rounds of mitotic divisions. Fill colors correspond to those used in (D), and the thickness of the outlines and connecting lines correspond to distance from the oocyte (thicker lines are closer to the oocyte, and thinner lines are further from the oocyte).

One of the most well-established examples of subcellular scaling is that of the nucleus. Cells typically maintain a constant ratio of nuclear to cytoplasmic volume, which appears to be actively maintained in healthy cells. When cells divide, or if cell size is artificially manipulated, the nucleus will adjust its size to re-establish the proper ratio (Balachandra et al., 2022; Cantwell and Nurse, 2019). Within the developing egg chamber, the growth of the nurse cells, their nuclei, and nucleoli is similar, which leads to isometric scaling of these structures (Diegmiller et al., 2021; Imran Alsous et al., 2017). Similar isometric scaling of nuclear, nucleolar, and cell size within the developing *C. elegans* intestine has also been observed (Uppaluri et al., 2016). Nurse cell nuclear growth is primarily driven by multiple rounds of endoreplication, and increased ploidy correlates with increased cell size (Diegmiller et al., 2021; Doherty et al., 2021; Orr-Weaver, 2015; Øvrebø and Edgar, 2018). Although all nurse cells and their nuclei grow during oogenesis, it has recently been demonstrated that nurse cell size is not uniform within the germline cell cluster; instead, there is a size hierarchy based on distance from the oocyte. The four posterior nurse cells directly connected to the oocyte are the largest, whereas the anterior most nurse cell that is four ring canals away from the oocyte is the smallest (Diegmiller et al., 2021; Imran Alsous et al., 2017). Therefore, within this system, we have the opportunity to monitor subcellular scaling in a growing tissue with a cell size gradient.

The germline ring canals are essential structures whose size must be tightly regulated to produce a viable mature egg. The germline cell cluster is formed through four rounds of mitosis; at the end of each division, the midbody is modeled to form the ring canal (Price et al., 2023). These mitotic divisions are synchronous across the cluster, which gives rise to a stereotypical pattern of connections (Fig. 1E). After completion of the four mitotic divisions, the ring canals begin to expand (Ong and Tan, 2010), ultimately reaching a final diameter of nearly 10 µm. Although it is known that ring canal growth is required for fertility, the scaling of ring canal size during oogenesis has not been determined.

Here, we characterize the scaling of two types of ring canals: the four ring canals that connect the posterior most nurse cells to the oocyte and the anterior most ring canal. We hypothesized that ring canal size and growth would correlate with that of the associated nurse cells; if that were true, we would have expected to see an anterior-posterior gradient in ring canal size and scaling. Instead, our data supported an alternative hypothesis that ring canal scaling is most closely linked to lineage. The largest, “first born” ring canals expand more slowly compared to smaller, later born ring canals. This scaling pattern is maintained even if polarized transport is reduced, ring canal size is altered, or in egg chambers with twice as many germline cells. Finally, analysis of a set of artificially selected “big egg” and “small egg” lines revealed that the lineage dependent pattern of scaling is maintained, but that subtle changes to ring canal scaling could provide a mechanism to regulate final egg size.

## Materials and Methods

### Fly Stocks and Maintenance

Flies were maintained on a cornmeal molasses diet at 18-25°C. The “small egg” (9.31.4, 8.26.2, 8.11.1, 7.17.4, and 9.2.2) and “big egg” (1.16.1, 2.15.4, 2.49.3, 1.40.2, and 3.34.1) *D. melanogaster* lines were obtained from C. Miles (Augustana University)(Jha et al., 2015; Miles et al., 2011), the *otu-GAL4,UAS-Dcr-2* (recombinant of BDSC #58424 and BDSC #24648) was provided by J. Merkle (University of Evansville), and the *UASp-ovhts:GFP* line was provided by L. Cooley (Gerdes et al., 2020). The following lines were obtained from the Bloomington *Drosophila* stock center: *w^1118^* (BDSC# 3605), *UAS-dhc64C-RNAi* (BDSC #36583), *matαTub-GAL4* (BDSC #7063), maternal triple driver MTD-GAL4 (BDSC #31777), *nos-GAL4* (BDSC #32563), and *UAS-CG34200/tom6-RNAi* (BDSC #44463).

### Immunofluorescence and Imaging

Female flies were maintained with ground yeast in the presence of males for ∼44-60 hours prior to dissection. Ovaries were dissected in Schneider’s media, fixed in 4% formaldehyde, and stained with an HtsRC antibody (1:20, hts RC DSHB), phalloidin (1:500; either TRITC or FITC conjugated; ECM Biosciences), and DAPI (1:500). Anti-mouse secondary antibodies were used at a 1:200 dilution (Jackson Immunolab). Tissue was imaged using a Nikon Ti2-E Inverted microscope equipped with a Yokogawa CSU-X1 Spinning Disk and Hamamatsu ORCA Fusion camera with a 20x PLAN APO/0.75 NA objective. Although we tried to be consistent in scaling confocal images in the figures, there were differences in fluorescence intensity within z-stacks, so scaling was adjusted to best highlight the structures of interest in each image.

### Mature Egg Collection and Volume Measurements

Male and female flies were incubated with ground yeast for approximately 24 hours at 25°C prior to transferring them to cages with apple juice plates and wet yeast. Mature eggs were collected and imaged using a Zeiss LP-520 and ProgRes MF camera (Jenoptik) using ProgRes Capture Pro Software (Jenoptik). Length and width measurements were made using Fiji (Schindelin et al., 2012), and volumes were calculated using the formula: Volume = 1/6*π*Width^2^*Length

### Ring canal and Nuclear Measurements

Nuclear and ring canal measurements were made using Fiji (Schindelin et al., 2012) with either the freehand drawing tool or the line tool respectively. Measurements were performed in the plane that contained the maximum cross section or diameter within the DAPI or HtsRC channel. For Fig. 3, egg chambers with low levels of ovhts::GFP expression were excluded from the analysis. Egg chamber volumes were calculated using the formula: Volume = 1/6*π*Width^2^*Length

### Statistical Analysis

Graphing and statistical analyses were performed using Prism 10.1.0 (GraphPad Software, LLC). Data are typically shown as individual data points or as the mean +/- s.d. or the 95% confidence interval as indicated in legends. For linear regression analysis, ANCOVA was used to determine whether there were significant differences in the slopes or y-intercepts. One-way ANOVA with Tukey’s multiple comparisons test was used to compare mature egg size (Fig. 5A). The sample size for each experiment (n) is indicated in the figure legends.

## Results

### Ring canal scaling is related to lineage

It has recently been shown that there is a size hierarchy within the germline cell cluster; the largest nurse cell nuclei are located at the posterior, whereas the smallest nucleus is within the anterior most nurse cell (Diegmiller et al., 2021; Imran Alsous et al., 2017). Recent work in this system has demonstrated that nurse cell nuclear growth is tightly coupled to cell growth (Diegmiller et al., 2021). Therefore, we can use nuclear size as a proxy for nurse cell size.

We first wanted to determine whether there were differences in the growth rate of nurse cell nuclei in different regions of the germline cell cluster. We focused our analysis on two sets of nurse cells – the four posterior nurse cells that are directly connected to the oocyte and the anterior most nurse cell (Fig. 1E). This should allow us to compare the relative growth rates of the largest and smallest nurse cells. We found that as the egg chamber increases in volume, the anterior and posterior nurse cell nuclei also increase in size, but at slightly different rates (Fig. 1A-B). If we plot these measurements on a log-log scale, then the slope of the regression line provides us with the scaling exponent, or the way that the size of one structure is related to the size of another structure. Although the intercept does not represent a biologically relevant measurement (since you cannot measure the size of one structure when the size of the other structure is 0), significant differences in the y-intercepts allow us to compare the relative size of structures between different conditions or genetic backgrounds. Because we are plotting the log of egg chamber volume and the log of nuclear area, isometric nuclear scaling would be reflected by a slope of 2/3 or 0.666; a smaller slope would indicate hypoallometric scaling, and a larger slope would indicate hyperallometric scaling. We found that the posterior nuclei scale nearly isometrically with egg chamber volume (Fig. 1B; scaling exponent of 0.652, 95% confidence interval is 0.627-0.678), which is consistent with what has been previously shown (Diegmiller et al., 2021). The nucleus in the anterior most nurse cell scales slightly hypoallometrically (Fig. 1B; scaling exponent of 0.626, 95% confidence interval of 0.593-0.659), but this difference is not significant (p>0.05). The y-intercepts of these two lines are significantly different (Fig. 1B; p<0.0001), which is consistent with the smaller size of the anterior cells (and their nuclei) that has been previously reported (Diegmiller et al., 2021; Imran Alsous et al., 2017). This suggests that in addition to absolute differences in nuclear size, there could be modest differences in nurse cell (and nuclear) growth rates based on position within the germline cell cluster.

If we hypothesize that ring canal growth correlates with nurse cell size and growth, then we would predict that the posterior ring canals should be larger and expand faster than the anterior ring canal. We measured the diameter of the ring canals that connect the same nurse cells to either the oocyte (for the posterior ring canals) or to another nurse cell (for the anterior ring canal). There was no difference in the scaling exponent between these two ring canal types (Fig. 1C; 95% confidence interval of 0.218-0.241 for the posterior ring canals and 0.209-0.250 for the anterior ring canals), but there was again a significant difference in the y-intercepts (Fig. 1C; p<0.0001), which reflects the smaller size of the anterior ring canals. At first glance, these data suggest ring canal size varies along the anterior-posterior axis, which correlates with nuclear size, but that ring canal scaling is similar across the germline cell cluster.

However, what was obvious from our analysis was that within an egg chamber, the size of the four posterior ring canals varied significantly (Fig. 1A,C). This variation in size could be explained by the emergence of one ring canal at the end of each of the four mitotic divisions that form the germline cyst. It has been shown that the ring canals formed during the earlier mitotic divisions are larger than those that are formed during the later divisions (Ong and Tan, 2010), at least within the germarium. Therefore, we hypothesized that lineage could be the source of the posterior ring canal size differences. If we separate out the posterior ring canals based on their mitotic division of origin (Fig. 1E; M1 for the first mitotic division, M2 for the second mitotic division, etc.), we found that the “first born” M1 ring canals were nearly always the largest, whereas the “last born” M4 ring canals were typically the smallest (Fig. 1A,D). In addition to these size differences, the posterior ring canals had different scaling exponents; the M1 ring canals had a significantly lower scaling exponent compared to the M2, M3, and M4 ring canals (M1 ring canals: 0.202, 95% confidence interval 0.188-0.216; M4 ring canals: 0.252, 95% confidence interval 0.234-0.271). Interestingly, the scaling exponent for the anterior ring canals, which are also formed during the fourth mitotic division, was not significantly different from that of the posterior M4 ring canals (Fig. 1D). This observation suggests that ring canal size and growth are not linked to nurse cell size, since the anterior nurse cells are the smallest and grow more slowly than the posterior nurse cells. Instead, our data support the hypothesis that ring canals scale differently based on their lineage. These differences could reflect a need for the initially smaller M4 ring canals to grow more quickly to try to “catch up” to the size of the larger M1 ring canals.

### Reducing directed transport reduces ring canal size and leads to a modest increase in scaling

Ring canals serve a specific purpose within the egg chamber by providing a channel for the directed transport of material from the nurse cells into the oocyte. Based on the observed size differences, we hypothesized that the larger M1 ring canals may allow more material to flow through them than the smaller M4 ring canals; this difference in flow could provide a mechanism to drive faster growth of the M4 ring canals. If the amount of directed transport through the ring canal inversely correlates with ring canal growth, then reducing transport should increase growth. It has been shown that the amount of biased transport of materials through the germline ring canals correlates with their position along the anterior-posterior axis (Lu et al., 2021). Movement of materials through the anterior ring canals is bidirectional, whereas the movement through the posterior ring canals is significantly biased toward the oocyte. Depletion of either dynein heavy chain (Dhc64C) or the linker protein Short stop (Shot) significantly reduces biased transport at the posterior, in turn reducing oocyte size (Lu et al., 2021). Therefore, if ring canal size scaling is sensitive to transport through the posterior ring canals, then depletion of Dhc64C should alter the scaling exponents.

We first confirmed that depletion of Dhc64C led to a significant reduction in oocyte size. To perform the depletion, we used *otu-GAL4,UAS-Dcr-2* which bypasses the requirement for dynein during earlier stages of oogenesis (Lu et al., 2022; McGrail and Hays, 1997). This phenotype was most obvious during mid-oogenesis (stage ∼5-8), so we focused our analysis on these egg chambers (Fig. 2A-B). We performed the same analysis of ring canal diameter in the control and *dhc64C-RNAi* egg chambers. We found that depletion of Dhc64C led to a significant decrease in ring canal diameter (Fig. 2C,S1A; y-intercepts for both ring canal types were significantly different from controls), and there was a modest, but not significant increase in ring canal scaling (Fig. 2C,S1A). A similar reduction in ring canal size was observed in Shot-depleted egg chambers (Lu et al., 2021). If we separated the ring canals based on lineage, we again found that depletion of Dhc64C led to a significant decrease in ring canal size and a consistent, but not significant increase in the scaling exponent for all ring canal types (Fig. 2D, S1B significant difference in y-intercepts for control/*dhc64C-RNAi* comparisons for M2, M3, M4 and the anterior ring canals, p<0.05). Despite these changes, the pattern of lineage-based size scaling was maintained.

**Figure 2:**
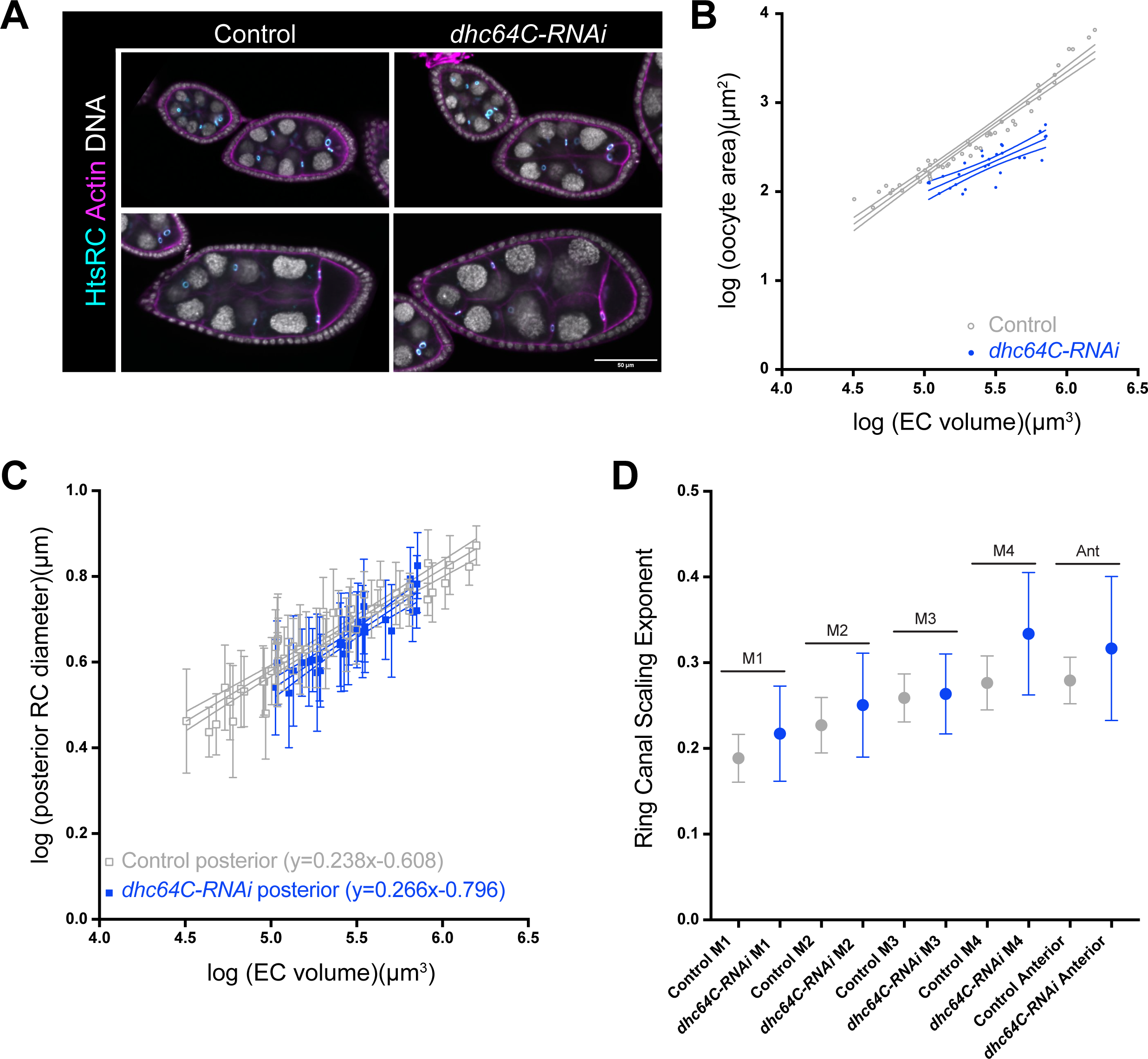
Reducing directed transport and oocyte growth has a modest effect on ring canal size and scaling. (A) Single plane confocal images of control and *dhc64C-RNAi* egg chambers (*otu-GAL4,UAS-Dcr-2*) stained with an HtsRC antibody, phalloidin, and DAPI. Scale bar is 50 µm. (B,C) log-log plots of egg chamber (EC) volume, oocyte area, or ring canal (RC) diameter from stage 5-10b. 95% confidence intervals are shown. Error bars in C are s.d. (D) Scaling exponent for ring canals separated by lineage. Error bars are 95% confidence intervals. n=49 egg chambers for control and 28 egg chambers for *dhc64C-RNAi*.

### Increasing ring canal size does not affect the lineage-based patterns, but it does lead to modest differences in scaling

If ring canal size inversely correlates with scaling such that larger ring canals expand more slowly than smaller ring canals, then increasing ring canal diameter should reduce the scaling exponent. To test this hypothesis, we over-expressed a GFP-tagged form of ovhts, which is cleaved to produce HtsRC::GFP (Gerdes et al., 2020). As was previously reported (Gerdes et al., 2020), we observed a consistent increase in size for all ring canal types (Fig. 3A-F; y-intercepts were significantly different for M1, M2, M4, and anterior RCs, p<0.0001), but the lineage-based pattern of scaling exponents was still observed (Fig. 3B-F). Although the scaling exponents were consistently lower in egg chambers over-expressing HtsRC::GFP, these differences were only significant for the M3 ring canals (Fig. 3D, p=0.008). This suggests that increasing ring canal size is not sufficient to significantly reduce scaling for all ring canal types.

**Figure 3:**
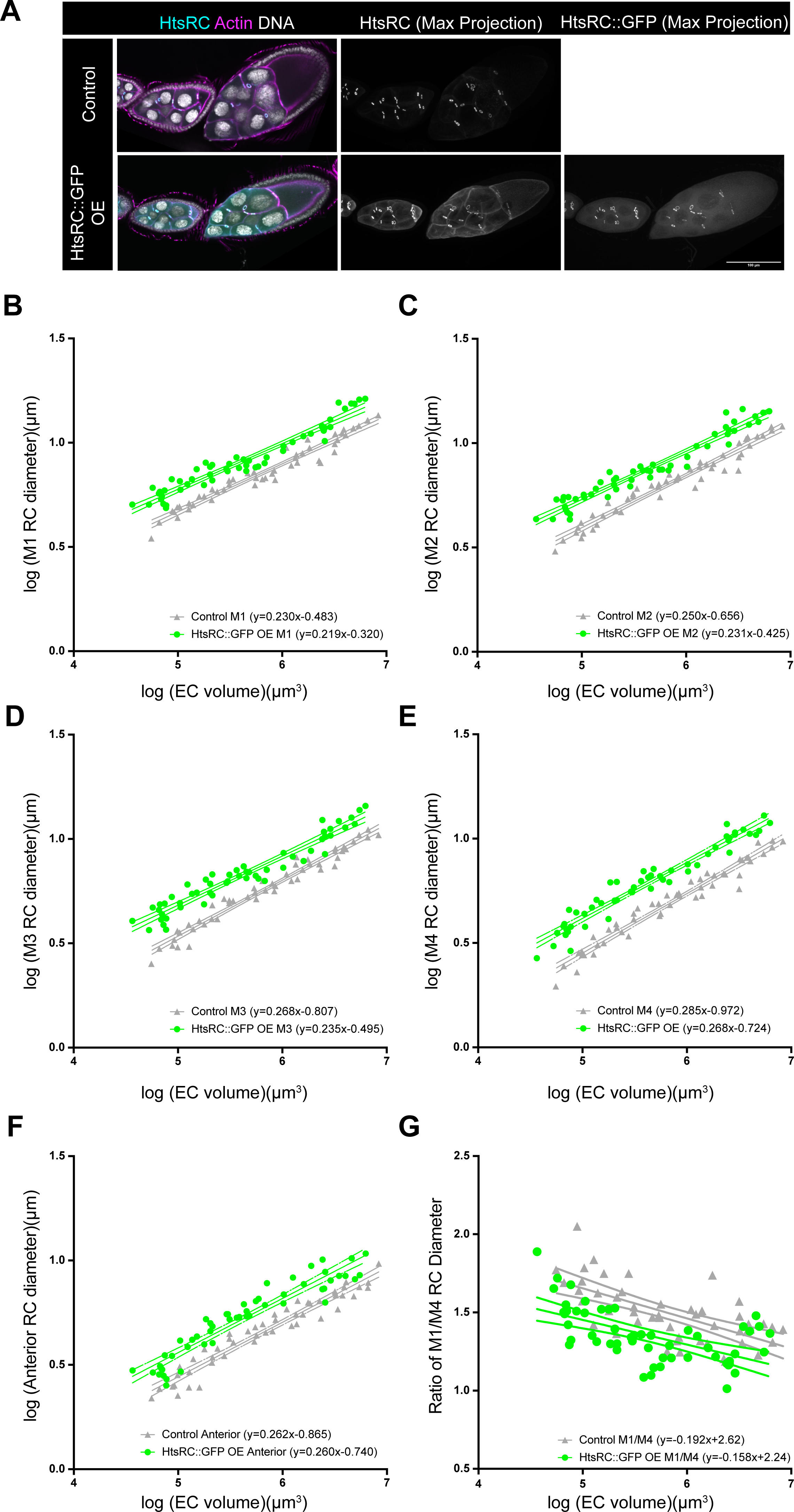
Over-expression of HtsRC::GFP increases ring canal size without affecting the lineage-based scaling. (A) Single plane or maximum intensity projections of confocal images of control and HtsRC::GFP over-expressing (OE) egg chambers (*matαTub-GAL4*) stained with an HtsRC antibody, phalloidin, and DAPI. Scale bar is 100 µm. (B-F) log-log plots of egg chamber (EC) volume and ring canal (RC) diameter and (G) the ratio of M1/M4 ring canal diameter from stage 5-10b. 95% confidence intervals are shown. n=50 egg chambers for control and 54 egg chambers for HtsRC::GFP OE.

Although the lineage-based scaling pattern in the HtsRC::GFP over-expressing egg chambers were similar to the control, our impression was that the differences in posterior ring canal size were not as large. Therefore, we calculated the ratio of the M1 ring canal diameter to the M4 ring canal diameter. In control egg chambers, this ratio is initially higher, but as the egg chamber increases in volume, the ratio decreases, approaching, but never reaching 1.0 (Fig. 3G). For the HtsRC::GFP over-expressing egg chambers, we still observed the same negative correlation, but the ratios were significantly lower than in controls (Fig. 3G; y-intercepts were significantly different, p<0.0001). This suggests that even though over-expression of HtsRC::GFP increases the size of all ring canals, the “last born” ring canals still grow faster and are sometimes even able to catch up to the size of the “first born” ring canals.

### Lineage-based ring canal scaling is maintained in egg chambers with twice as many germline cells

Thus far, we have described previously unappreciated trends in the size scaling relationships of the germline ring canals based on lineage. Although size may contribute to the growth differences, lineage seems to be the most consistent predictor of scaling. To further test this relationship, we turned to a condition in which egg chambers contain twice as many germline cells. As part of another project, it was found that when a shRNA construct was used to induce a depletion of CG34200, or Tom6, using *nanos-GAL4*, many egg chambers contained 31 nurse cells and ring canals, with a single oocyte connected to five posterior nurse cells (Fig. 4A). This phenotype could occur if the germline cells go through an additional (fifth) mitotic division; alternatively, if there is a delay in abscission following the germline stem cell division, this could give rise to a 2-cell cystoblast, which then goes through four mitotic divisions to produce the 32-cell cyst (Mathieu et al., 2013; Matias et al., 2015). Additional work to further characterize the Tom6 depletion phenotype is in progress (manuscript in preparation), but we could use this condition to determine whether ring canal scaling changes in egg chambers that contain additional germline cells and ring canals.

**Figure 4:**
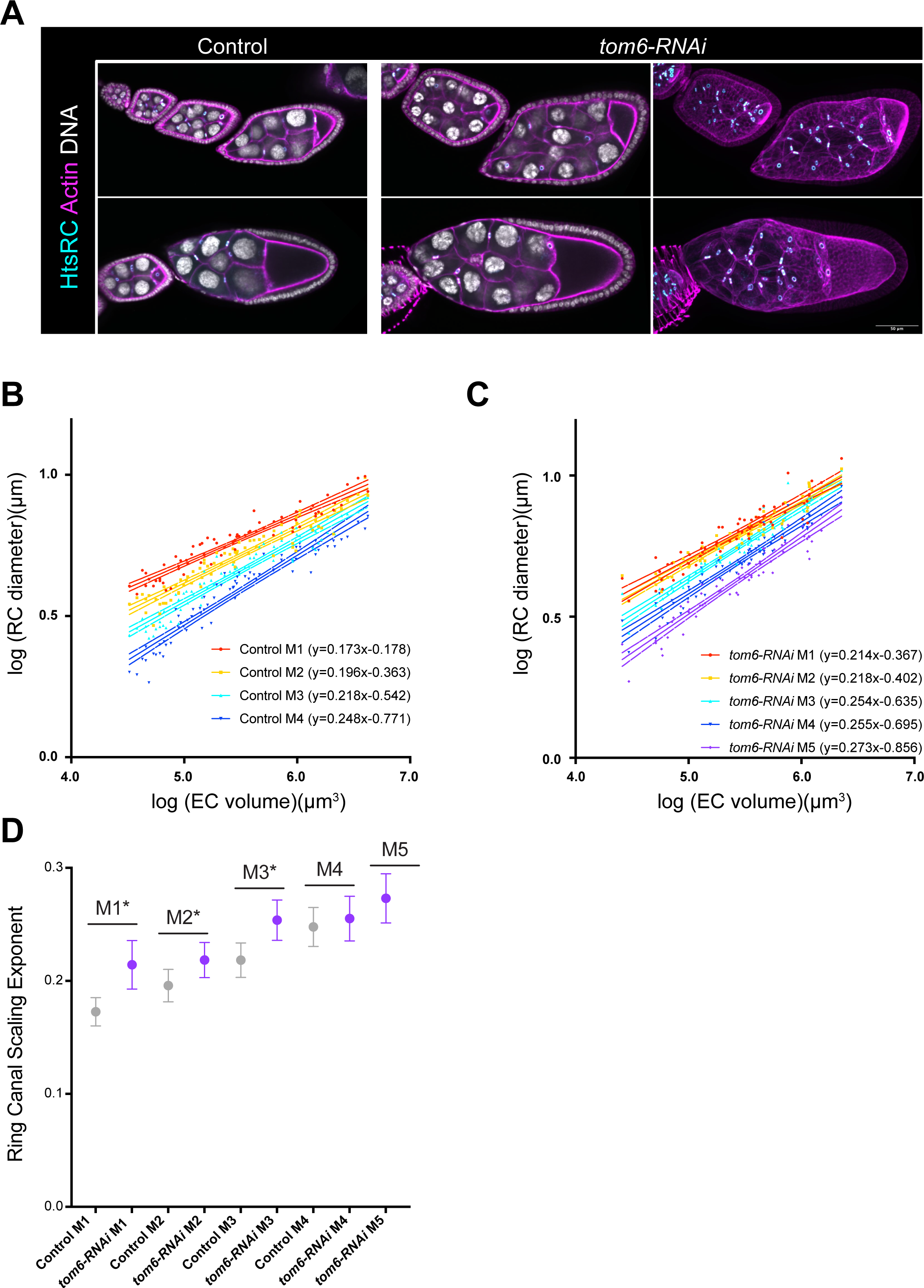
Depletion of Tom6 increases the number of germline cells but does not alter the pattern of ring canal scaling. (A) Single plane confocal images of control and *tom6-RNAi* egg chambers (*nos-GAL4*) stained with an HtsRC antibody, phalloidin, and DAPI. Maximum intensity projections of the *tom6-RNAi* egg chambers are included to show the additional nurse cells and ring canals in these egg chambers. Scale bar is 50 µm. (B,C) log-log plots of egg chamber (EC) volume and ring canal (RC) diameter from stage 5-10b. n=74 egg chambers for control and 64 egg chambers for *tom6-RNAi*. 95% confidence intervals are shown. (D) Scaling exponent for ring canals separated by lineage. Error bars indicate 95% confidence interval. Asterisks indicate significant difference between control and *tom6-RNAi* (ANCOVA, p<0.05).

When Tom6 was depleted from the germline, we saw similar trends for ring canal scaling. The ring canal scaling exponent increased from M1 to M5 (Fig. 4B-D). Although the ring canals in *tom6-RNAi* egg chambers were typically larger, some ring canal types had a significantly larger scaling exponent (Fig. 4D) compared to controls. Therefore, although size could impact scaling, our data again support the model that lineage is the strongest predictor of ring canal scaling.

### Scaling differences were observed across a collection of “big” and “small” egg lines

When considering the importance of size scaling within the germline of the developing egg chamber, we wondered whether it would be maintained in egg chambers that develop into eggs of different sizes or whether differences in ring canal size and/or scaling could contribute to differences in final egg volume. We chose to monitor scaling within a collection of *D. melanogaster* lines that were artificially selected to produce either “big” or “small” eggs (Jha et al., 2015; Miles et al., 2011). We first confirmed that these lines still produce significantly larger or smaller mature eggs; all the “big egg” lines produced significantly larger mature eggs than all the “small egg” lines (Fig. 5A; p<0.0001 ANOVA with Tukey’s post-hoc). From this collection, we performed further analysis on two of the “big egg” lines (1.40.2 and 3.34.1) and two of the “small egg” lines (7.17.4 and 9.31.4).

**Figure 5:**
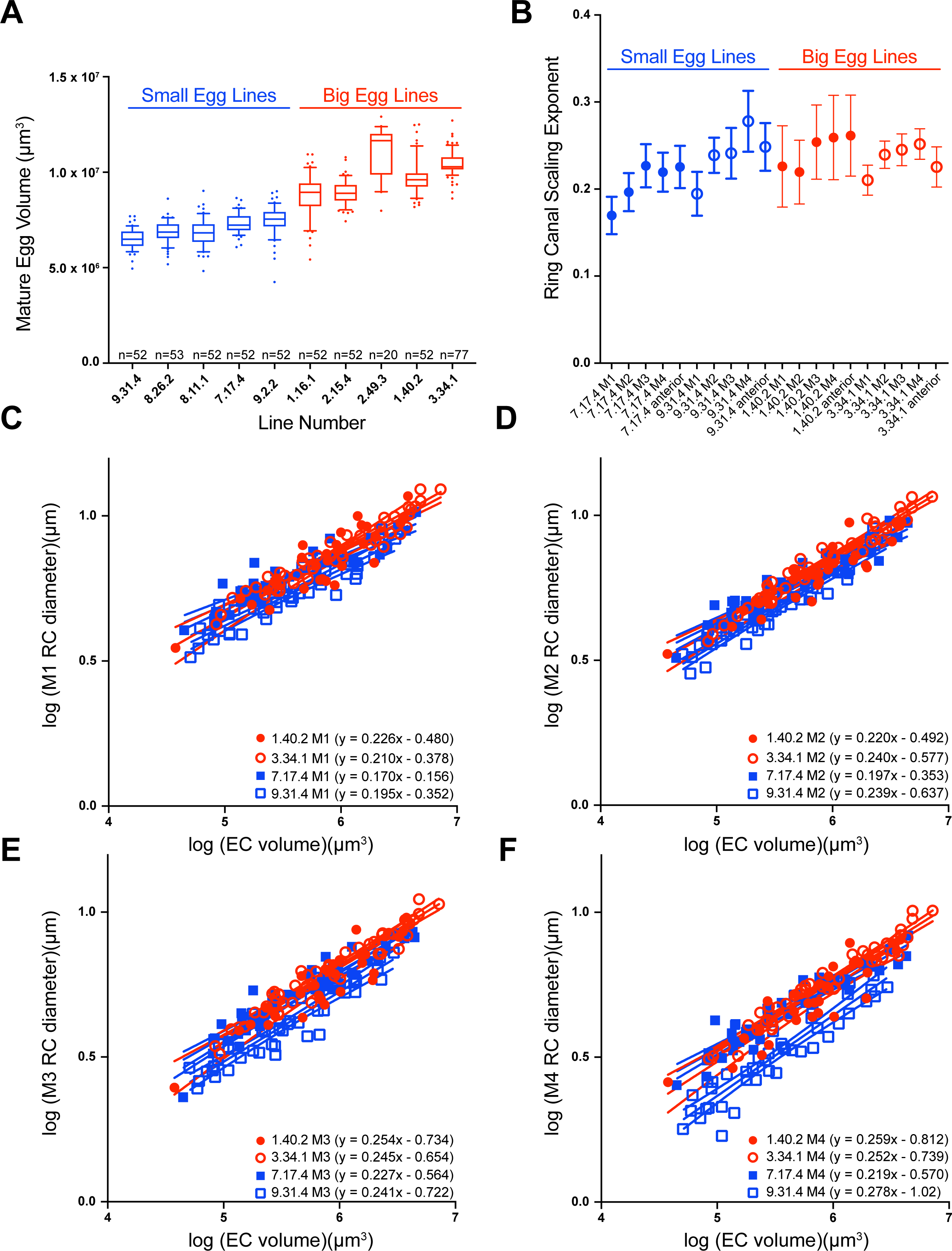
Differences in posterior ring canal size and scaling in “big egg” and “small egg” *D. melanogaster* lines. (A) Box and whiskers plot (10-90^th^ percentile) of mature egg volume of “big egg” and “small egg” *D. melanogaster* lines. One-way ANOVA with Tukey’s multiple comparisons test indicates mature eggs from all “small egg” lines (9.31.4, 8.26.2, 8.11.1, 7.17.4, and 9.2.2) are significantly smaller than all “big egg” lines (1.16.1, 2.15.4, 2.49.3, 1.40.2, and 3.34.1) (p<0.0001). n= 20-77 eggs per line. (B) Scaling exponent for ring canals separated by lineage. Error bars indicate 95% confidence interval. There were significant differences in the scaling exponent across the four lines for M1, M2, and M4 (ANCOVA, p<0.05). (C-F) log-log plots of egg chamber (EC) volume and ring canal (RC) diameter in two “big egg” (1.40.2 and 3.34.1) and two “small egg” (7.17.4 and 9.31.4) lines separated by lineage. 95% confidence intervals are shown. n=29 egg chambers for 1.40.2, 46 egg chambers for 3.34.1, 44 egg chambers for 7.17.4, and 43 egg chambers for 9.31.4.

In this set of four lines, we went on to monitor ring canal size. We hypothesized that if lineage determines ring canal scaling, then we should see the same patterns across the “big egg” and “small egg” lines, where the M1 ring canals have a lower scaling exponent than the M4 ring canals. When comparing ring canal size and scaling between the “big egg” and “small egg” lines, we considered different possibilities. For example, if the M1 ring canals are larger in the “big egg” lines than the M1 ring canals in the “small egg” lines, then the scaling could be similar between those lines, which would support the model that lineage, and not ring canal size, most reliably determines scaling. If ring canal size does impact growth, then in this scenario, the M1 ring canals in the “big egg” lines might have a lower scaling exponent than the M1 ring canals in the “small egg” lines. Alternatively, ring canal size may be similar across the “big egg” and “small egg” lines, but differences in scaling could provide a mechanism to alter final egg size.

After completing ring canal measurements, we found that the lineage-based pattern of scaling exponents was consistent with what we had observed previously. Within each line, the scaling exponents typically increased from M1 to M4 (Fig. 5B-F), and there was no significant difference in the scaling exponent for the M4 ring canal compared to the anterior ring canal (Fig. 5B). We were surprised to see that there was not a consistent relationship between ring canal size and final egg volume. Although the “big egg” lines did tend to contain larger ring canals, that was not always the case. However, the line that produces the smallest mature eggs, 9.31.4, did consistently have smaller ring canals than the other lines (Fig. 5A,C-F; y-intercept for 9.31.4 was significantly different than “big egg” lines for all ring canal types; p<0.0001); this difference was most obvious when comparing the size of the M4 ring canals (Fig. 5F). Although the ring canals were smaller in the 9.31.4 line, the scaling exponents were not significantly different than those of the two “big egg” lines (Fig. 5B-F). This suggests that reducing ring canal size without altering scaling could provide a mechanism to reduce final egg size.

The other “small egg” line, 7.17.4, showed a different behavior. This line consistently had scaling exponents for the posterior ring canals that were significantly smaller than the “big egg” lines (Fig. 5B-F); however, the anterior ring canal scaling was not different (Fig. S2A). Unlike the other “small egg” line, the size of the posterior ring canals in 7.17.4 was more similar to the size of those in the “big egg” lines, but the lower scaling exponent suggests that they may be expanding at a slower rate. This line also had a consistently lower nuclear scaling exponent for all nurse cell types (Fig. S2B). This suggests that coordinated changes in ring canal and nurse cell (nuclear) scaling could contribute to changes in final egg size. Additional work will be necessary to determine whether there are other size and/or scaling differences across this collection of “big egg” and “small egg” lines, which could provide insight into different developmental strategies that could be used to alter final egg size.

## Discussion

We still have much to learn about how cells within developing tissues coordinate their size and the size and number of their subcellular structures, but here, we have characterized size scaling of the germline ring canals within the developing egg chamber. Our data support the model that birth order inversely correlates with ring canal scaling; the larger, “first born” ring canals have a lower scaling exponent than the smaller ring canals derived from later mitotic divisions (Fig. 1-4). Although some data support a modest effect of directed transport (Fig. 2) or ring canal size (Fig. 3), lineage consistently provided the strongest predictor of ring canal scaling. Analysis of artificially selected lines that produce eggs of different sizes (Fig. 5) suggests that within *D. melanogaster*, there could be multiple strategies used to alter final egg size.

### Ring Canal Scaling

Ring canals are known to undergo significant growth during oogenesis, but to our knowledge, this is the first time that their scaling has been described. It was not surprising that the diameter of the ring canals would increase more slowly than other structures within the egg chamber, such as the nurse cell nuclei or nucleoli (Diegmiller et al., 2021). Although they are a common feature of developing germline cells across the animal kingdom, most intercellular bridges are much smaller (typically less than 2 µm in diameter) than the germline ring canals within the egg chamber (Haglund et al., 2011). A recent study reported that when they are formed, the ratio of intercellular bridge diameter to cell diameter was ∼5-15% (Singh et al., 2022), which suggests that there might be a physical limit to the size of a stable intercellular bridge. One way to overcome this limitation might be to alter ring canal structure. Most other intercellular bridges contain a single “rim,” but the germline ring canals within the egg chamber, which undergo a nearly 20-fold increase in diameter, contain two (Haglund et al., 2011). Perhaps this more complex structure enables more growth than other intercellular bridges or stabilizes these larger channels.

Ring canals initially differ in size due to differences in the degree of contractile ring constriction at the end of each mitosis, producing smaller ring canals with each round of division (Ong 2010). It has been shown that once the mitotic divisions are complete, the M1 ring canal is the first to initiate the growth phase (Ong 2010), and we assumed that growth would be similar for all ring canal types. Instead, our data support a lineage-based effect on ring canal scaling, with the ring canals from earlier mitotic divisions increasing in size more slowly than the ring canals from later divisions (Fig. 1-5). The scaling differences could reflect nurse cell size differences; if that were the case, then the posterior M4 ring canal would scale differently than the anterior ring canal, since the anterior cells (and their nuclei) are always smaller than the posterior cells (Fig. 1) (Diegmiller et al., 2021; Imran Alsous et al., 2017). The observation that the scaling exponents for the M4 posterior ring canal and the anterior ring canal were not significantly different (Fig. 1) argues against that model.

Another possibility is that there could be structural differences in the ring canals that are formed during the different mitotic divisions. Early work suggests that the germline cells do not double in size between each round of mitosis (Koch and King, 1966), so as the cluster develops, the dividing cells are smaller and smaller. In the *C. elegans* embryo, the contractile ring contains structural memory of cell size; the contractile rings in larger cells divide faster than those in smaller cells. This structural memory has been proposed to be due to the incorporation of different numbers of contractile units into rings of different sizes (Carvalho et al., 2009). Because the ring canal develops from the remodeled midbody, and the midbodies become smaller with each germline cell division (Price et al., 2023), it is possible that the structure of the nascent intercellular bridge may retain some memory of the size of the dividing cell, which may then impact its subsequent rate of growth.

Additional structural differences could also explain lineage-based growth differences. For example, if the M1 contractile ring constricts less than contractile rings formed in subsequent divisions, perhaps this larger intercellular bridge must be reinforced by the recruitment of more actin filaments or stabilized by more cell-cell junctions, which may impact growth. Extensive cell-cell contacts have been observed in the developing germline cell cysts (Loyer et al., 2015; Riparbelli et al., 2022; Tilney et al., 1996), but a comparison of their distribution within different regions of the cluster has not been performed.

If ring canal size correlates with oocyte-directed transport, then lineage-based ring canal scaling could be used to ultimately achieve similar rates of transport through all ring canals. If this were the case, then we would expect to see ring canals of different lineages approach a similar final size; the initially smaller M4 ring canals grow faster to catch up to the size of the larger M1 ring canals. When we compared the ratio of the M1 diameter to the M4 diameter, we found that this ratio did approach, but never reached 1.0. When the ring canals were larger in the HtsRC::GFP over-expressing egg chambers, this ratio was significantly lower (Fig. 3G), but we still observed a lineage-based effect on scaling. It is possible that the ring canals do not adjust their growth rate based on the size of other ring canals, or it could be that because our over-expression begins around stage 2-3 of oogenesis, lineage-based structural differences have already been established. If there is an upper limit or a stage-specific optimal size for transport of materials into the oocyte, then perhaps the larger ring canals hit that upper limit or reach the optimal size sooner, which could explain the reduced scaling exponents when HtsRC is over-expressed (Fig. 3). Additional work will be necessary to further explore the relative rate of transport through different types of ring canals and whether that changes as the ring canals increase in size.

One final explanation we considered is that the growth of a posterior ring canal could depend on how many other ring canals exist within that germline cell. For example, the M1 ring canal that we were monitoring is connecting the oocyte to a nurse cell that contains three additional ring canals (M2, M3, and M4 types), whereas the posterior M4 ring canal is the only intercellular bridge for that nurse cell (Fig. 1E). Therefore, the differences in scaling could be due to the presence or absence of additional ring canals within that cell. One argument against that model is that in Tom6-depleted egg chambers that contain twice as many germline cells, each of the posterior nurse cells had an additional ring canal, but the scaling exponents were not lower than what we observed in the controls (Fig. 4). Therefore, additional work is necessary to determine the molecular mechanisms underlying these scaling differences.

Although lineage does appear to be the strongest predictor of ring canal scaling, we cannot rule out a contribution of ring canal size to scaling. In egg chambers depleted of Dhc64C, there was a reduction in ring canal size and increase in scaling (Fig. 2), and in egg chambers over-expressing HtsRC::GFP, there was a consistent increase in ring canal size and reduction in scaling (Fig. 3). However, the *tom6-RNAi* egg chambers typically contained larger ring canals, but some ring canal types showed higher scaling exponents (Fig. 4). Therefore, analysis of additional conditions will be necessary to determine the impact of ring canal size on scaling.

### The lineage-based pattern of ring canal scaling is observed in lines that produce eggs of different sizes

Final egg size is likely determined by many genes that impact the size or growth of multiple structures during oogenesis. One possibility is that when producing eggs of different sizes, the size of all structures is similarly altered. For example, when generating larger mature eggs, flies produce larger egg chambers that contain larger ring canals that expand at a specific rate based on their lineage. For three of the lines analyzed (1.40.2, 3.34.1, and 9.31.4), we found that there were no significant differences in the lineage-based ring canal scaling exponents. In these egg chambers, the line that produced the smallest mature eggs (9.31.4) tended to have the smallest ring canals relative to egg chamber volume (Fig. 5C-F), which was especially obvious for the M4 ring canals (Fig. 5F). Analysis of these lines, therefore, is consistent with this developmental strategy.

Another approach could be to increase or decrease ring canal scaling, which could then affect the transfer of materials into the oocyte, ultimately changing final egg size. Whole genome sequencing of the entire collection of artificially selected “big egg” and “small egg” lines identified a set of candidate genes that may contribute to egg size evolution (Jha et al., 2015); that list included many proteins that localize to the germline ring canals or have been implicated in ring canal size regulation, including Cheerio/Filamin, Flapwing, Mbs, and Short stop (Jha et al., 2015). Cheerio/Filamin localizes to germline ring canals and is required for formation of the actin-rich inner rim (Robinson et al., 1997). Flapwing and Mbs have been shown to work together to inhibit myosin activity during incomplete cytokinesis, and mutants form smaller ring canals (Ong et al., 2010; Yamamoto et al., 2013). As previously discussed, Short stop is a linker protein that promotes directed transport into the oocyte (Lu et al., 2021). One of the small egg lines (7.17.4) had a lower scaling exponent for all posterior ring canals (Fig. 5), which suggests that this could be a mechanism to alter final egg size.

Many studies have demonstrated the impact of environmental conditions on egg size, but the molecular mechanisms that control this life history trait are still not known. Some insight has come from a recent study that found that honey bees could reversibly alter egg size in response to the perceived colony size. The larger eggs that were produced in response to a smaller perceived colony size expressed higher levels of the GTPase, Rho1, and other related cytoskeletal proteins; depletion of Rho1 reduced honey bee final egg size (Han et al., 2022). In *Drosophila,* Rho family members have been implicated in regulating germline actin structures, including the ring canals (Murphy and Montell, 1996). Additional studies will be necessary to confirm the connection between ring canal scaling and final egg size and to further explore pathways that may be involved in establishing or adjusting ring canal scaling. Our study focused on ring canal growth prior to nurse cell dumping, so it remains to be determined how this stage of oogenesis proceeds in the “big egg” and “small egg” lines and whether there could be differences in the duration of the dumping phase and/or the degree of completion of material transfer between the different lines. In addition, because our study looked at a narrow range of final egg sizes, future work could expand to include analysis of oogenesis in insects that produce much larger and much smaller eggs.

## Acknowledgements

We would like to thank Seth Donoghue and Andrew Stoehr for helpful suggestions on the manuscript and Cecelia Miles for the big and small egg lines. This work was supported by the National Institutes of Health (NIH R15HD084243 to L.L. and R15GM099054 to W.G.). The confocal microscope was purchased with support from a Major Research Instrumentation (MRI) award from the National Science Foundation (#2116348). Stocks obtained from the Bloomington Drosophila Stock Center (NIH P40OD018537) were used in this study. The following antibody was obtained from the Developmental Studies Hybridoma Bank, created by the NICHD of the NIH and maintained at the University of Iowa, Department of Biology, Iowa City, IA 52242: hts RC antibody developed by L. Cooley.

## References

Ali-Murthy, Z., Fetter, R. D., Wang, W., Yang, B., Royer, L. A. and Kornberg, T. B. (2021). Elimination of nurse cell nuclei that shuttle into oocytes during oogenesis. J. Cell Biol. 220, e202012101.

Balachandra, S., Sarkar, S. and Amodeo, A. A. (2022). The Nuclear-to-Cytoplasmic Ratio: Coupling DNA Content to Cell Size, Cell Cycle, and Biosynthetic Capacity. Annu. Rev. Genet. 56, 165–185.

Cantwell, H. and Nurse, P. (2019). Unravelling nuclear size control. Curr. Genet. 65, 1281–1285.

Carvalho, A., Desai, A. and Oegema, K. (2009). Structural Memory in the Contractile Ring Makes the Duration of Cytokinesis Independent of Cell Size. Cell 137, 926– 937.

Diegmiller, R., Doherty, C. A., Stern, T., Alsous, J. I. and Shvartsman, S. Y. (2021). Size scaling in collective cell growth. Development dev.199663.

Doherty, C. A., Diegmiller, R., Kapasiawala, M., Gavis, E. R. and Shvartsman, S. Y. (2021). Coupled oscillators coordinate collective germline growth. Dev. Cell 56, 860–870.e8.

Gayon, J. (2000). History of the Concept of Allometry1. Am. Zool. 40, 748–758.

Gerdes, J. A., Mannix, K. M., Hudson, A. M. and Cooley, L. (2020). HtsRC-Mediated Accumulation of F-Actin Regulates Ring Canal Size During Drosophila melanogaster Oogenesis. Genetics 216, 717–734.

Haglund, K., Nezis, I. P. and Stenmark, H. (2011). Structure and functions of stable intercellular bridges formed by incomplete cytokinesis during development. Commun. Integr. Biol. 4, 1–9.

Han, B., Wei, Q., Amiri, E., Hu, H., Meng, L., Strand, M. K., Tarpy, D. R., Xu, S., Li, J. and Rueppell, O. (2022). The molecular basis of socially induced egg-size plasticity in honey bees. eLife 11, e80499.

Huxley, J. S. and Teissier, G. (1936). Terminology of Relative Growth. Nature 137, 780–781.

Imran Alsous, J., Villoutreix, P., Berezhkovskii, A. M. and Shvartsman, S. Y. (2017). Collective Growth in a Small Cell Network. Curr. Biol. 27, 2670–2676.e4.

Januschke, J., Gervais, L., Dass, S., Kaltschmidt, J. A., Lopez-Schier, H., Johnston, D. S., Brand, A. H., Roth, S. and Guichet, A. (2002). Polar Transport in the Drosophila Oocyte Requires Dynein and Kinesin I Cooperation. Curr. Biol. 12, 1971–1981.

Jha, A. R., Miles, C. M., Lippert, N. R., Brown, C. D., White, K. P. and Kreitman, M. (2015). Whole-Genome Resequencing of Experimental Populations Reveals Polygenic Basis of Egg-Size Variation in *Drosophila melanogaster*. Mol. Biol. Evol. 32, 2616–2632.

Koch, E. A. and King, R. C. (1966). The origin and early differentiation of the egg chamber of Drosophila melanogaster. J. Morphol. 119, 283–303.

Loyer, N., Kolotuev, I., Pinot, M. and Le Borgne, R. (2015). *Drosophila* E-cadherin is required for the maintenance of ring canals anchoring to mechanically withstand tissue growth. Proc. Natl. Acad. Sci. 112, 12717–12722.

Lu, W., Lakonishok, M. and Gelfand, V. I. (2021). Gatekeeper function for Short stop at the ring canals of the Drosophila ovary. Curr. Biol. 31, 3207–3220.e4.

Lu, W., Lakonishok, M., Serpinskaya, A. S. and Gelfand, V. I. (2022). A novel mechanism of bulk cytoplasmic transport by cortical dynein in Drosophila ovary. eLife 11, e75538.

Marshall, W. F. (2020). Scaling of subcellular structures. Annu. Rev. Cell Dev. Biol. 36, 219–236.

Mathieu, J., Cauvin, C., Moch, C., Radford, S. J., Sampaio, P., Perdigoto, C. N., Schweisguth, F., Bardin, A. J., Sunkel, C. E., McKim, K., et al. (2013). Aurora B and Cyclin B Have Opposite Effects on the Timing of Cytokinesis Abscission in *Drosophila* Germ Cells and in Vertebrate Somatic Cells. Dev. Cell 26, 250–265.

Matias, N. R., Mathieu, J. and Huynh, J.-R. (2015). Abscission Is Regulated by the ESCRT-III Protein Shrub in Drosophila Germline Stem Cells. PLOS Genet. 11, e1004653.

McGrail, M. and Hays, T. S. (1997). The microtubule motor cytoplasmic dynein is required for spindle orientation during germline cell divisions and oocyte differentiation in Drosophila. Development 124, 2409–2419.

Miles, C. M., Lott, S. E., Hendriks, C. L. L., Ludwig, M. Z., Manu, null, Williams, C. L. and Kreitman, M. (2011). Artificial selection on egg size perturbs early pattern formation in Drosophila melanogaster. Evol. Int. J. Org. Evol. 65, 33–42.

Murphy, A. M. and Montell, D. J. (1996). Cell type-specific roles for Cdc42, Rac, and RhoL in Drosophila oogenesis. J. Cell Biol. 133, 617–630.

Ong, S. and Tan, C. (2010). Germline cyst formation and incomplete cytokinesis during Drosophila melanogaster oogenesis. Dev. Biol. 337, 84–98.

Ong, S., Foote, C. and Tan, C. (2010). Mutations of DMYPT cause over constriction of contractile rings and ring canals during Drosophila germline cyst formation. Dev. Biol. 346, 161–169.

Orr-Weaver, T. L. (2015). When bigger is better: the role of polyploidy in organogenesis. Trends Genet. TIG 31, 307–315.

Øvrebø, J. I. and Edgar, B. A. (2018). Polyploidy in tissue homeostasis and regeneration. Dev. Camb. Engl. 145, dev156034.

Price, K. L., Tharakan, D. M. and Cooley, L. (2023). Evolutionarily conserved midbody remodeling precedes ring canal formation during gametogenesis. Dev. Cell 58, 474–488.e5.

Riparbelli, M. G., Persico, V. and Callaini, G. (2022). Cell-to-Cell Interactions during Early Drosophila Oogenesis: An Ultrastructural Analysis. Cells 11, 2658.

Robinson, D. N., Smith-Leiker, T. A., Sokol, N. S., Hudson, A. M. and Cooley, L. (1997). Formation of the Drosophila Ovarian Ring Canal Inner Rim Depends on Cheerio. Genetics 145, 1063–1072.

Schindelin, J., Arganda-Carreras, I., Frise, E., Kaynig, V., Longair, M., Pietzsch, T., Preibisch, S., Rueden, C., Saalfeld, S., Schmid, B., et al. (2012). Fiji - an Open Source platform for biological image analysis. Nat. Methods 9, 10.1038/nmeth.2019.

Singh, J., Imran Alsous, J., Garikipati, K. and Shvartsman, S. Y. (2022). Mechanics of stabilized intercellular bridges. Biophys. J. 121, 3162–3171.

Tilney, L. G., Tilney, M. S. and Guild, G. M. (1996). Formation of actin filament bundles in the ring canals of developing Drosophila follicles. J. Cell Biol. 133, 61– 74.

Uppaluri, S., Weber, S. C. and Brangwynne, C. P. (2016). Hierarchical Size Scaling during Multicellular Growth and Development. Cell Rep. 17, 345–352.

Yamamoto, S., Bayat, V., Bellen, H. J. and Tan, C. (2013). Protein Phosphatase 1ß Limits Ring Canal Constriction during Drosophila Germline Cyst Formation. PLOS ONE 8, e70502.

