## Supplemental Figures and legends for "Lineage-based scaling of germline intercellular bridges during oogenesis"

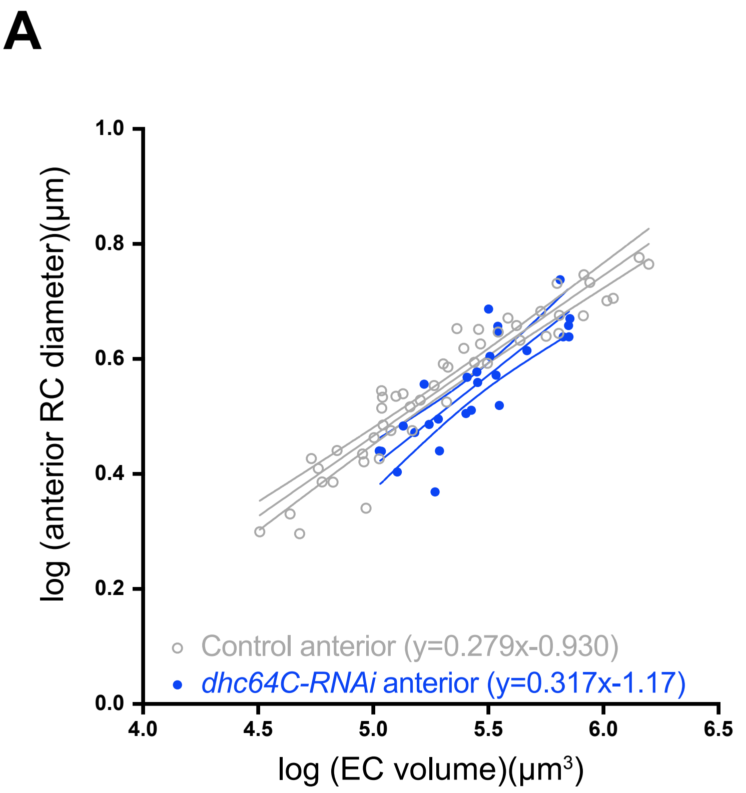
A
1.0
0.8
0.6
0.4
0.2
0.0
log (anterior RC diameter)(μm)
4.0
4.5
5.0
5.5
6.0
6.5
log (EC volume)(μm³)
Control anterior (y=0.279x-0.930)
dhc64C-RNAi anterior (y=0.317x-1.17)

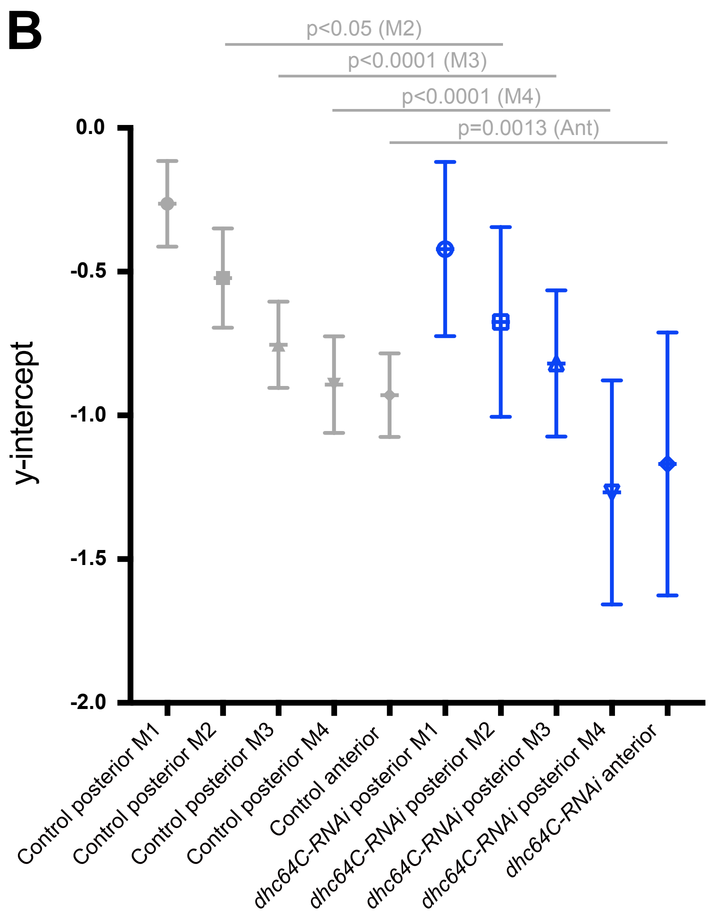
B
p<0.05 (M2)
p<0.0001 (M3)
p<0.0001 (M4)
p=0.0013 (Ant)
0.0
-0.5
-1.0
-1.5
-2.0
y-intercept
Control posterior M1
Control posterior M2
Control posterior M3
Control posterior M4
Control anterior
dhc64C-RNAi posterior M1
dhc64C-RNAi posterior M2
dhc64C-RNAi posterior M3
dhc64C-RNAi posterior M4
dhc64C-RNAi anterior

Figure S2

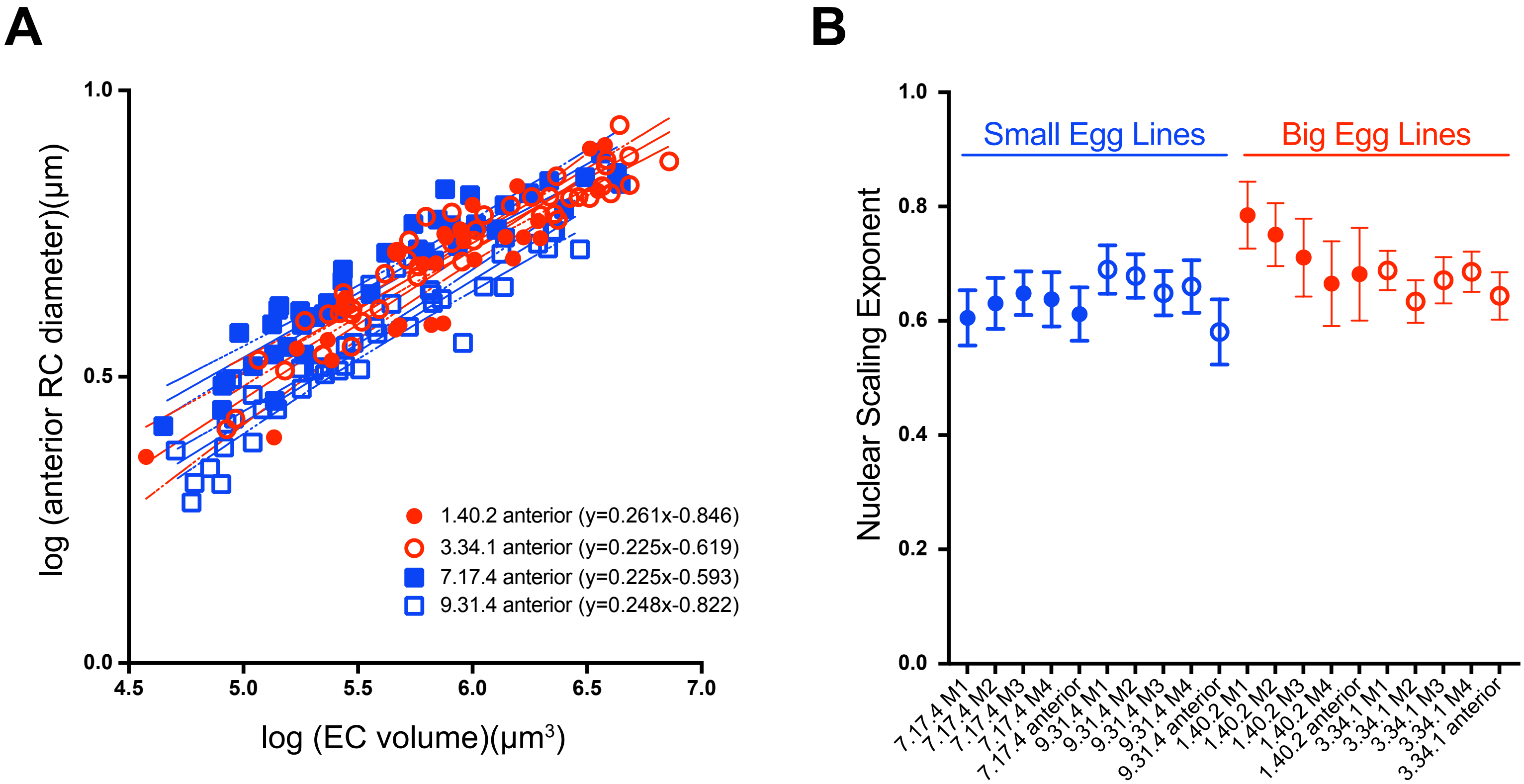
A
1.0
0.5
0.0
4.5
5.0
5.5
6.0
6.5
7.0
log (anterior RC diameter)(µm)
log (EC volume)(µm³)
1.40.2 anterior (y=0.261x-0.846)
3.34.1 anterior (y=0.225x-0.619)
7.17.4 anterior (y=0.225x-0.593)
9.31.4 anterior (y=0.248x-0.822)
B
Nuclear Scaling Exponent
1.0
0.8
0.6
0.4
0.2
0.0
Small Egg Lines
Big Egg Lines
7.17.4 M1
7.17.4 M2
7.17.4 M3
7.17.4 M4
7.17.4 anterior
9.31.4 M1
9.31.4 M2
9.31.4 M3
9.31.4 M4
9.31.4 anterior
1.40.2 M1
1.40.2 M2
1.40.2 M3
1.40.2 M4
1.40.2 anterior
3.34.1 M1
3.34.1 M2
3.34.1 M3
3.34.1 M4
3.34.1 anterior

**Figure S1: Reducing oocyte growth has a modest effect on ring canal size and scaling.** (A) log-log plot of egg chamber (EC) volume and anterior ring canal (RC) diameter from stage 5-10b. 95% confidence intervals are shown. (B) y-intercepts of regression lines from log-log plots of egg chamber volume and ring canal diameter (related to Fig. 2). Error bars are 95% confidence intervals. n=49 egg chambers for control and 28 egg chambers for *dhc64C-RNAi*.

**Figure S2: Anterior ring canal and nuclear scaling in "big egg" and "small egg" *D. melanogaster* lines.** (A) log-log plot of egg chamber (EC) volume and anterior ring canal (RC) diameter in two big egg (1.40.2 and 3.34.1) and two small egg (7.17.4 and 9.31.4) lines. 95% confidence intervals are shown. (B) Scaling exponent (+/- 95% confidence interval) for nuclei based on lineage. n=29 egg chambers for 1.40.2, 46 egg chambers for 3.34.1, 44 egg chambers for 7.17.4, and 43 egg chambers for 9.31.4.
